# Timing of TORC1 inhibition dictates Pol III involvement in longevity in *Caenorhabditis elegans*

**DOI:** 10.1101/2022.10.17.512483

**Authors:** Yasir Malik, Isabel Goncalves Silva, Rene Rivera Diazgranados, Colin Selman, Nazif Alic, Jennifer M. A. Tullet

**Affiliations:** Division of Natural Sciences, University of Kent, Canterbury, Kent, CT2 7NZ; Institute of Healthy Ageing, UCL Department of Genetics, Evolution & Environment, London WC1E 6BT; Institute of Biodiversity, Animal Health and Comparative Medicine, University of Glasgow, Glasgow, G12 8QQ

**Keywords:** RNA Polymerase III, mTOR, TORC1, longevity, ageing, *Caenorhabditis elegans*

## Abstract

Organismal growth and lifespan are inextricably linked. Target of Rapamycin (TOR) signalling regulates protein production for growth and development, but if reduced extends lifespan across species. Reduction of the enzyme RNA polymerase III, which transcribes tRNAs and 5S rRNA, also extends longevity. Here, we identify a temporal genetic relationship between TOR and Pol III in *C. elegans*, showing that they collaborate to regulate progeny production and lifespan. Interestingly, the lifespan interaction between Pol III and TOR is only revealed when TOR signaling is reduced specifically in adulthood demonstrating the importance of timing to control TOR regulated developmental *vs* adult programs. Additionally, we show that Pol III acts in *C. elegans* muscle to promote both longevity and healthspan and that reducing Pol III even in late adulthood is sufficient to extend lifespan. This demonstrates the importance of Pol III for lifespan and age-related health in adult *C. elegans*.

## Introduction

In eukaryotic cells, three multi-subunit RNA polymerase (Pol) enzymes, Pol I, II, III transcribe the nuclear genome. Each polymerase synthesises a distinct set of genes: Pol II transcribes all coding genes to generate mRNA while polymerases Pol I and III only transcribe non-coding genes^1^. RNA Pol III is comprised of 17 subunits and is specialised for the transcription of short, abundant, non-coding RNA transcripts. Its main gene products are transfer RNAs (tRNAs) and 5S ribosomal RNA (rRNA), which are abundantly transcribed and required for protein synthesis^2^. Other Pol III transcripts include small nuclear (snRNAs) and small nucleolar (snoRNAs) both of which are implicated in RNA processing, the 7SK snRNAs that plays role in regulating transcription, as well as other small RNAs including Y RNA, L RNA, vault RNA and U6 spliceosomal RNA^3,4^ Pol III-mediated transcription regulates a wide range of biological processes including cell and organismal growth^5–8^, the cell cycle^9^, differentiation^8,10,11^, development ^12^, regeneration^13^ and cellular responses to stress^14^. Pol III subunits have also been implicated in a wide variety of disease states and lifespan^15–17^.

Reducing activity of Pol III by partially downregulating its individual subunits can significantly extend lifespan in multiple model organisms including baker’s yeast *Saccharomyces cerevisiae*, the nematode *Caenorhabditis elegans* and fruit fly *Drosophila melanogaster*^18^. In *C. elegans, rpc-1* encodes the largest Pol III subunit and its downregulation by RNAi increases lifespan^18^. In *D. melanogaster* lifespan is increased in heterozygous females with a single functional copy of gene encoding for Pol III-specific subunit C53. Pol III activity in the fly gut limits survival and can drive ageing specifically from the gut stem-cell compartment. Importantly, in flies Pol III knockdown is downstream of the Target of Rapamycin pathway, an evolutionarily conserved regulator of longevity^18,19^

The Target of Rapamycin (TOR) kinase is a serine/threonine kinase that regulates growth and development chiefly by modulating protein synthesis in response to changes in nutrients and other cues^19^ (Fig 1A). TOR can be part of two complexes and it is specifically the TOR complex 1 (TORC1) that drives growth, development, and anabolic metabolism^19,20^. In *C. elegans*, complete loss of TOR/*let-363* is lethal, while depletion of TOR by RNAi or pharmacologically extends lifespan^21^ (Fig 1A).

**Figure 1.**
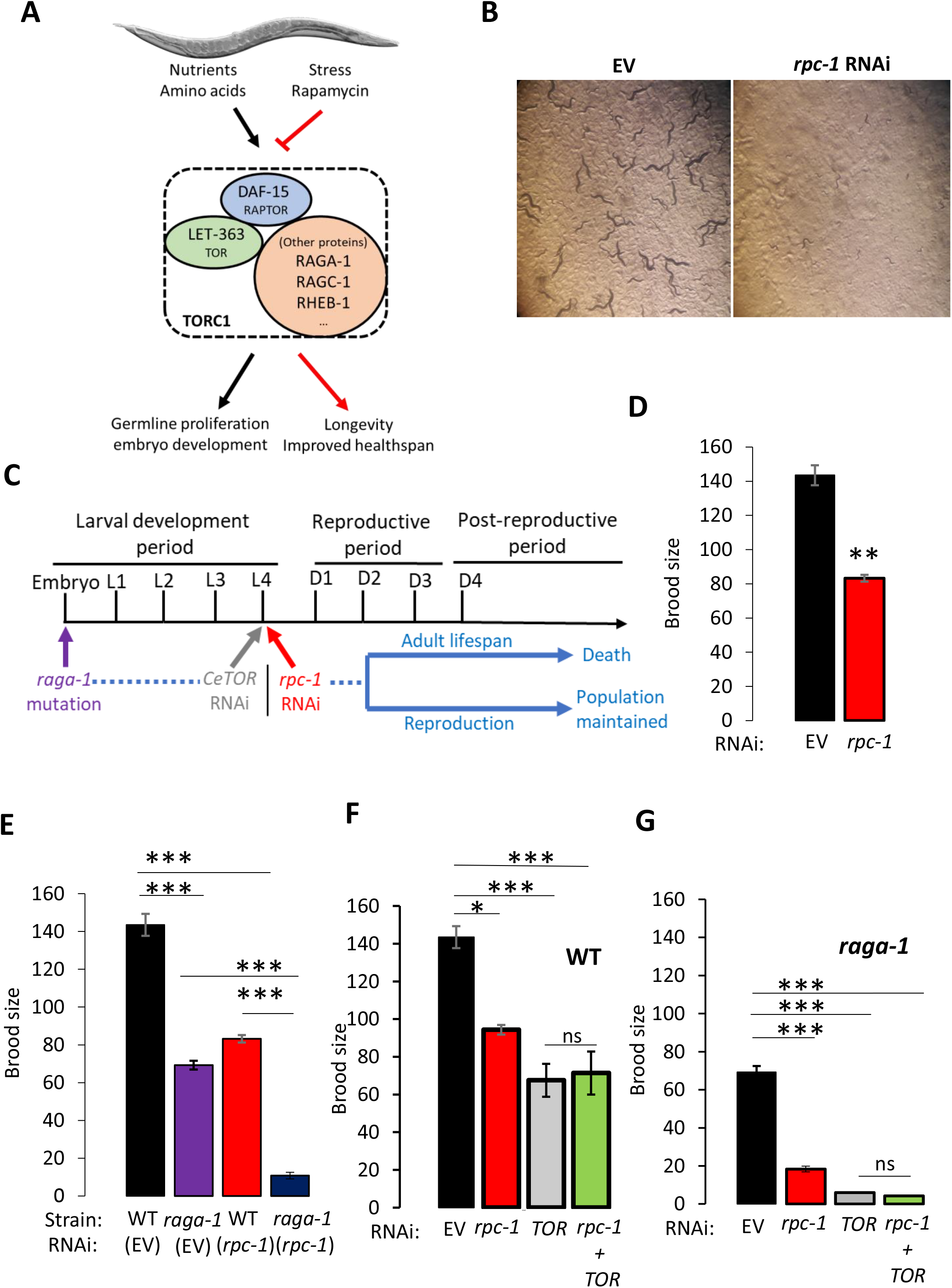
TOR signalling and RNA Polymerase III interact to control reproduction. **A)** Schematic showing the *C. elegans* TORC1 complex and its potential downstream outcomes in response to nutrient cues. **B)** *rpc-1* RNAi leads to L3 stage developmental arrest. This RNAi treatment typically leads to a 60% reduction in *rpc-1* mRNA levels ^18^. **C)** Schematic showing timings of the *rpc-1* and TOR interventions: for *rpc-1* and *CeTOR* RNAi worms were raised on OP50 and moved to the appropriate RNAi at the L4 stage; loss-of-function mutation in *raga-1(ok386)* is present throughout life, from the embryonic stage. **D - F)** Interaction between the TOR pathway and *rpc-1* for brood size. Combined data for three biological replicates is shown. EV, Empty Vector, One-way ANOVA: *,*p*<0.05,**; *p* <0.005; ***; *p*<0.0005; ns, non-significant.

Here we show that TOR and Pol III act together to control developmental and adult lifespan. Reducing TOR signaling from embryonic stages together with Pol III inhibition demonstrated their genetic interaction to control progeny production; but their ability to collaborate and regulate lifespan is only revealed when TOR signaling is reduced specifically in adulthood. This demonstrates the critical temporal importance of TOR signaling throughout life, how this alters it’s interactions with key transcriptional regulators, and the impact of this on developmental *vs* longevity programs. In addition, we show that Pol III acts in *C. elegans* muscle to promote both longevity and healthspan, and that late-life Pol III knockdown is sufficient to extend lifespan. Overall, our data uncover a mechanism which describes the interactions of TORC1 and Pol III in reproduction and longevity, and the importance of Pol III for lifespan and age-related health in adult *C. elegans*.

## Results

### RNA Polymerase III is important for *C. elegans* reproduction

RNA polymerase III is essential for growth and development^6,7,22^. Indeed, when we reduced Pol III mRNA expression by feeding *C. elegans* bacteria expressing a dsRNA for *rpc-1* from the embryonic stage, the worms exhibited growth defects and more than 90% of worms arrested at L2-L3 larval stage, whereas all animals on control RNAi developed to adults within 3 days (Fig. 1B). This confirms the requirement of Pol III for normal larval development in *C. elegans*. On the other hand, the requirement for Pol III in reproduction has not been examined. To determine the effect of Pol III knockdown on progeny production, we raised the worms to early adulthood on the standard laboratory food source *E. coli* OP50-1, transferring to *rpc-1* RNAi at the L4 larval stage (Fig. 1C). We found that *C. elegans* treated with *rpc-1* RNAi at this late larval stage developed into adults but displayed a dramatic reduction in their total brood size compared to control (∼ 40% *p*=2.98E-06, Fig.1D). This suggests that Pol III inhibition from late larval development is sufficient to reduce progeny production. Taken together, these data show that Pol III is required for normal organismal growth and reproductive processes.

### TOR signaling interacts genetically with RNA Polymerase III to control *C. elegans* reproduction

Reproduction and embryonic development in worms is an intense anabolic process requiring high levels of protein translation^23,24^. The TORC1 pathway acts as a master switch for governing cellular growth while Pol III transcribes two of the most important non-coding RNA families (tRNAs and rRNAs) required for protein synthesis^2^. To test whether Pol III could act downstream of TORC1 to control reproduction, we examined the epistatic relationship between *rpc-1* knockdown and *C. elegans* TORC1 using progeny production as a readout. In *C. elegans*, the gene encoding RagA (a RAS-related GTP-binding protein) acts as a positive regulator of the TORC1 kinase^25^ (Fig. 1A) and animals carrying the *raga-1(ok386)* mutation have constitutively low TORC1 signaling throughout development and adulthood^26^ (Fig. 1C). *raga-1* mutants develop normally but adults have a reduced brood size compared to WT (Fig. 1E, *p*=4.81E-06). However, treatment of *raga-1(ok386)* mutants with *rpc-1* RNAi resulted in a further reduction in progeny numbers compared to either *raga-1* mutation or *rpc-1* RNAi alone (Fig. 1E, *p*=7.91E-11 and *p*= 6.67E-8 respectively), implying that TOR and Pol III act independently to control progeny production.

In *C. elegans*, TORC1 is controlled by several inputs, and its inactivation can be achieved by several different genetic mechanisms^21^. We were concerned that the *raga-1(ok386)* mutation was not sufficient to completely reduce TORC1 signaling, so to address this, we depleted the *C. elegans* TOR kinase. Complete depletion of *C. elegans* TOR (*CeTOR)* is lethal so *CeTOR* RNAi was fed from the L4 stage (Fig. 1C). Similar to Pol III RNAi and *raga-1* mutation, *CeTOR* RNAi reduced brood size compared to control (Fig. 1F, *p*=4.9E-4), but combining *CeTOR* RNAi with *rpc-1* RNAi was non-additive compared to either *rpc-1* or TOR RNAi (Fig. 1F, *p*=0.57 and *p*= 0.62 respectively). In contrast to our initial result, this implies that Pol III and TOR act in the same pathway to regulate *C. elegans* reproduction. To explore this epistatic relationship further, and be sure that we fully reduced TORC1 signaling we examined progeny production in *raga-1(ok386)* mutants treated with *CeTOR* RNAi. Depletion of *CeTOR* from early adulthood resulted in a further reduction of *raga-1(ok386)* mutant brood size (Fig. 1G, *p*=1.31E-5) indicative that the initial *raga-1* mutation was incomplete. However, this was non-additive with *rpc-1* RNAi (Fig. 1G, *p*=0.80). Together, our data suggest that TORC1 and Pol III act in the same pathway to affect organismal reproduction.

### TOR and RNA Polymerase III interact genetically and temporally to control adult lifespan

TOR inhibition extends lifespan in a range of model organisms from yeast to mammals^27^. In *C. elegans*, depletion of TOR signaling either by *raga-1* loss-of-function mutation or *CeTOR* RNAi extends lifespan^26,28^. Given that Pol III and TORC1 act epistatically in respect to progeny production (Figs. 1E-G), we wanted to test the genetic interaction between TOR signaling and Pol III for *C. elegans* adult lifespan. As above, we reduced TOR signaling at two different time points: from embryonic stages using *raga-1* mutation and from the L4 stage using *CeTOR* RNAi, whilst consistently initiating Pol III knockdown at the L4 stage (Fig. 1C). We found that although *raga-1(ok386)* mutants are long-lived the addition of *rpc-1* RNAi further extended this lifespan indicating that TOR and Pol III are acting in parallel (Fig. 2A, Table 1). To rule out the possibility that *raga-1* mutation is incomplete and thus masking an epistatic relationship between TOR and Pol III we examined the simultaneous knock down of *raga-1* with *CeTOR* with and without *rpc-1* RNAi (Figs. 1A and 1C). In our hands *raga-1* mutation and *CeTOR* RNAi were not additive, however the addition of *rpc-1* RNAi did further increase the *raga-1; CeTOR* lifespan (Fig. 2B, Table 1). This supports our conclusion that TOR signaling engages additional, Pol III independent, mediators to control lifespan if knocked out during development (Fig. 2C, Table 1).

**Table 1.**

| Figure | Trial | Strain | Genotype | Mean lifespan (days) | Extension (%) *raga-1* | Extension (%) WT | p value (Log-rank) vs | N dead (total) |
| --- | --- | --- | --- | --- | --- | --- | --- | --- |
| 2C | 3 | WT | Control RNAi | 10.83 |  |  |  | 93 |
| 2C | 3 | WT | *rpc-1* RNAi | 11.72 |  | 8.22 | WT<0.001 | 91 |
| 2C | 3 | WT | *CeTOR* RNAi | 11.52 |  | 6.37 | WT<0.05<br>WT *rpc-1*:NS | 95 |
| 2B, 2C | 3 | WT | *rpc-1* RNAi dil | 11.73 |  | 8.31 | WT<0.001<br>WT *rpc-1*:NS<br>WT *ceTOR*:NS | 91 |
| 2B, 2C | 3 | WT | *CeTOR* RNAi dil | 11.84 |  | 9.33 | WT<0.001<br>WT *rpc-1*:NS<br>WT *ceTOR*:NS<br>WT *rpc-1*:NS | 72 |
| 2B, 2C | 3 | WT | *rpc-1* RNAi + *CeTOR* RNAi | 12.4 |  | 14.50 | WT<0.001<br>WT *rpc-1*<0.05<br>WT *ceTOR*<0.05<br>WT *rpc-1* :NS<br>WT *ceTOR* :NS | 84 |
| 2A | 3 | *raga-1(ok386)* | Control RNAi | 20.38 |  | 70.83 | WT<0.001<br>WT *rpc-1*<0.001<br>WT *ceTOR*<0.001<br>WT *rpc-1* <0.001<br>WT *ceTOR* <0.001<br>WT *rpc-1* + *CeTOR*<0.001 | 90 |
| 2A | 3 | *raga-1(ok386)* | *rpc-1* RNAi | 21.9 | 7.46 | 83.57 | WT<0.001<br>WT *rpc-1*<0.001<br>WT *ceTOR*<0.001<br>WT *rpc-1* <0.001<br>WT *ceTOR* <0.001<br>WT *rpc-1* + *CeTOR*<0.001<br>*raga-1*<0.05 | 81 |
| 2B | 3 | *raga-1(ok386)* | *CeTOR* RNAi | 22.03 | 8.10 | 84.66 | WT<0.001<br>WT *rpc-1*<0.001<br>WT *ceTOR*<0.001<br>WT *rpc-1* <0.001<br>WT *ceTOR* <0.001<br>WT *rpc-1* + *CeTOR*<0.001<br>*raga-1*<0.05<br>*raga-1* *rpc-1*:NS | 80 |
| 2B | 3 | *raga-1(ok386)* | *rpc-1* RNAi dil | 21.84 | 7.16 | 83.07 | WT<0.001<br>WT *rpc-1*<0.001<br>WT *ceTOR*<0.001<br>WT *rpc-1* <0.001<br>WT *ceTOR* <0.001<br>WT *rpc-1* + *CeTOR*<0.001<br>*raga-1*<0.05<br>*raga-1* *rpc-1*:NS<br>*raga-1* *ceTOR*:NS | 81 |

**Figure 2.**
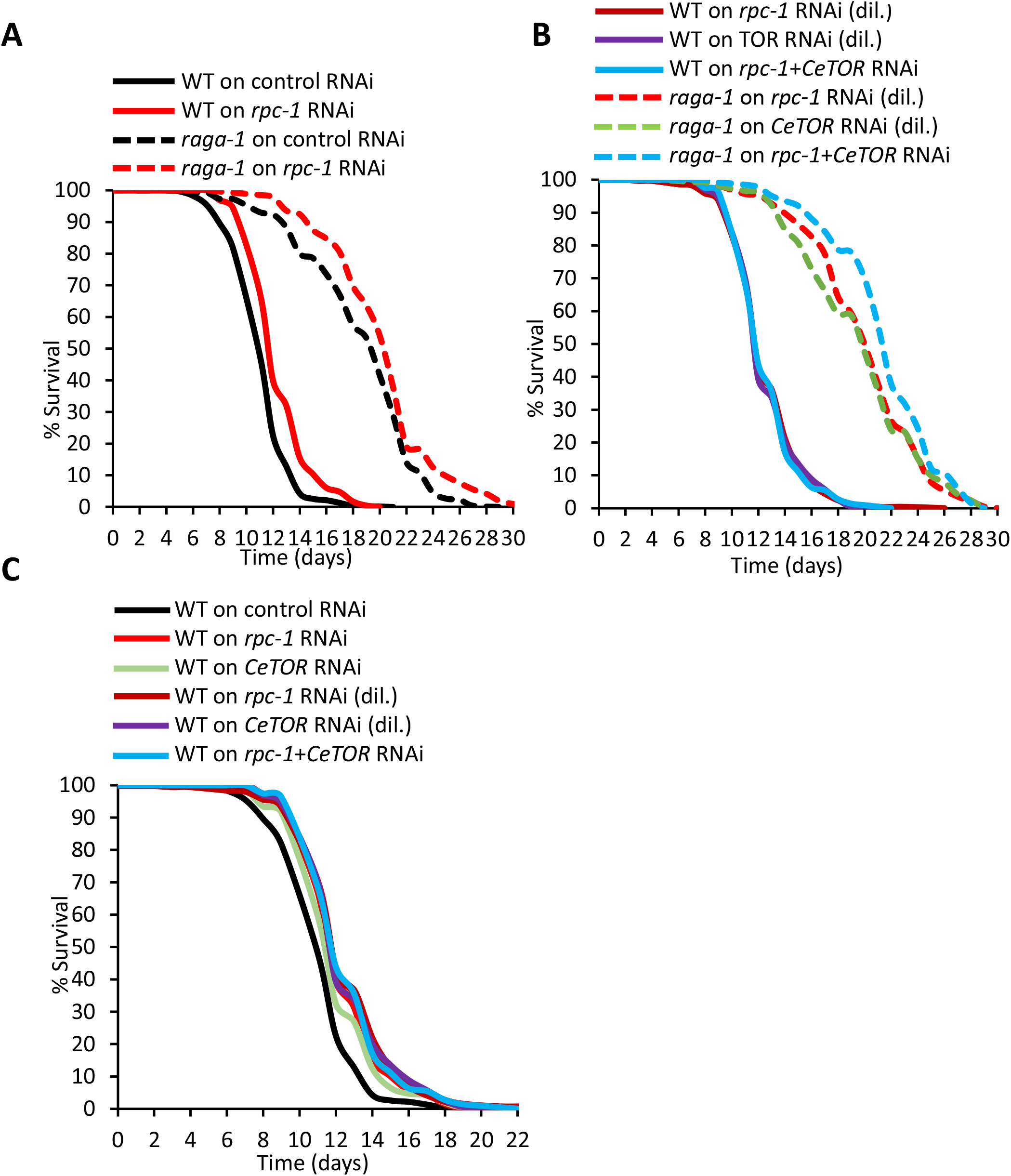
TOR signalling and RNA Polymerase III interact temporally to control lifespan. **A)** *rpc-1* RNAi and *raga-1(ok386)* additively increases lifespan. **B)** *CeTOR* RNAi and *raga-1(ok386)* mutation extend lifespan and are additive with *rpc-1* RNAi. **C)** *CeTOR* RNAi interacts with *rpc-1* RNAi to extend lifespan. **A-C)** One representative experiment is shown, refer to Table 1 for data on all replicates and statistical analysis.

Inhibition of TOR signaling in adult flies, using rapamycin, causes activation of RNA Pol III ^18^. In addition, the lifespan increases incurred by rapamycin treatment and Pol III knockdown are non-additive, indicating that Pol III mediates the effects of TOR for longevity^18^. To test whether a similar scenario exists in worms, we knocked down TOR signaling only from the L4 stage using *CeTOR* RNAi and examined its epistatic interaction with Pol III. Interestingly, although both knockdown of *CeTOR* or *rpc-1* from the L4 stage extended lifespan, there was no additive effect when the RNAi treatments were given simultaneously. This indicates that Pol III mediates the effects of TOR signaling in adulthood to control lifespan in *C. elegans* (Fig. 2C, Table 1). Taken together, our data suggest that the genetic interaction between TOR signaling and RNA Pol III is dependent on the timing of TOR reduction and that whilst reducing TOR signaling during development engages mediators in addition to Pol III to control lifespan, TOR reduction specifically in adulthood acts through Pol III to promote longevity.

### RNA Polymerase III does not interact with global protein synthesis, insulin or germline signalling to control lifespan

Reducing TOR signaling during development engages mediators distinct from Pol III to affect lifespan (Fig. 1C). To identify these, we tested the genetic interaction of Pol III with candidate mediators implicated in protein synthesis or developmental processes. The ribosomal S6 Kinase (S6K) is a direct target of the TOR kinase and regulates global translation^29,30^. In *C. elegans, rsks-1* encodes for S6K and animals lacking this gene are long-lived^31^ (Fig. S1A, Table S1). We found that this longevity was additive with that incurred by *rpc-1* RNAi treatment suggesting that Pol III and S6K do not interact to extend lifespan (Fig. S1A, Table S1). In mammals another TOR target is the translational repressor, eukaryotic initiation factor 4E-binding protein (4eBP) ^29,32^. *C. elegans* does not encode 4eBP, but animals with mutations in the translation initiation factors, *ife-2* /eIF4E and *ppp-1*/eIF2Bgamma, exhibit disruptions in protein translation^33^. However, although deletion of either *ife-2* or *ppp-1* increased *C. elegans* lifespan, this was additive with *rpc-1* RNAi in both instances (Fig. S1 B-C, Table S1). In addition to TOR signalling, lifespan is also controlled by other signaling pathways implicated in development and growth^34^. Mutation of the insulin receptor extends lifespan across species and has been implicated in protein synthesis and turnover in *C. elegans*^35,36^. In *C. elegans, daf-2* encodes the insulin receptor and *daf-2* loss-of-function mutants are long-lived^37^ (Fig. S2 A, Table S2). However, knockdown of Pol III using *rpc-1* RNAi from the L4 stage was additive with *daf-2* lifespan, suggesting that Pol III-does not interact with insulin signalling for lifespan. Loss of germline signalling also consistently increases lifespan in model organisms^38^. The *C. elegans* mutants *glp-1(e2141ts)* and *glp-4 (bn2)* are both sterile and live significantly longer than wild-types^39,40^. However, both of these mutations were additive with *rpc-1* RNAi (Fig. S2 B-C, Table S2) implying that that Pol III-inhibition induced longevity is independent of germline signaling. Taken together, these data show that Pol III does not interact with global translation mechanisms, insulin or germline signalling to control lifespan.

### RNA Polymerase III knockdown acts in specific tissues to extend *C. elegans* lifespan

In both *C. elegans* and *D. melanogaster* the longevity phenotype caused by Pol III knockdown appears to be mediated by the intestine, but in flies, neuronal Pol III may also contribute ^18^. The contribution of tissues other than the intestine has not been tested in worms, so to characterize the expression pattern of Pol III in *C. elegans*, we examined the expression of an *rpc-1::gfp::3xflag* translational reporter in wild type adult animals ^41^. We found RPC-1::GFP in a wide variety of tissues including nuclei of muscle, intestine, hypodermis, neurons and germline cells (Fig. 3A). Treatment of this reporter strain with *rpc-1* RNAi however reduced expression of RPC-1::GFP in all of these tissues (Fig. S3 A) indicating that these tissues could contribute to the Pol III-inhibition longevity phenotype.

**Figure 3.**
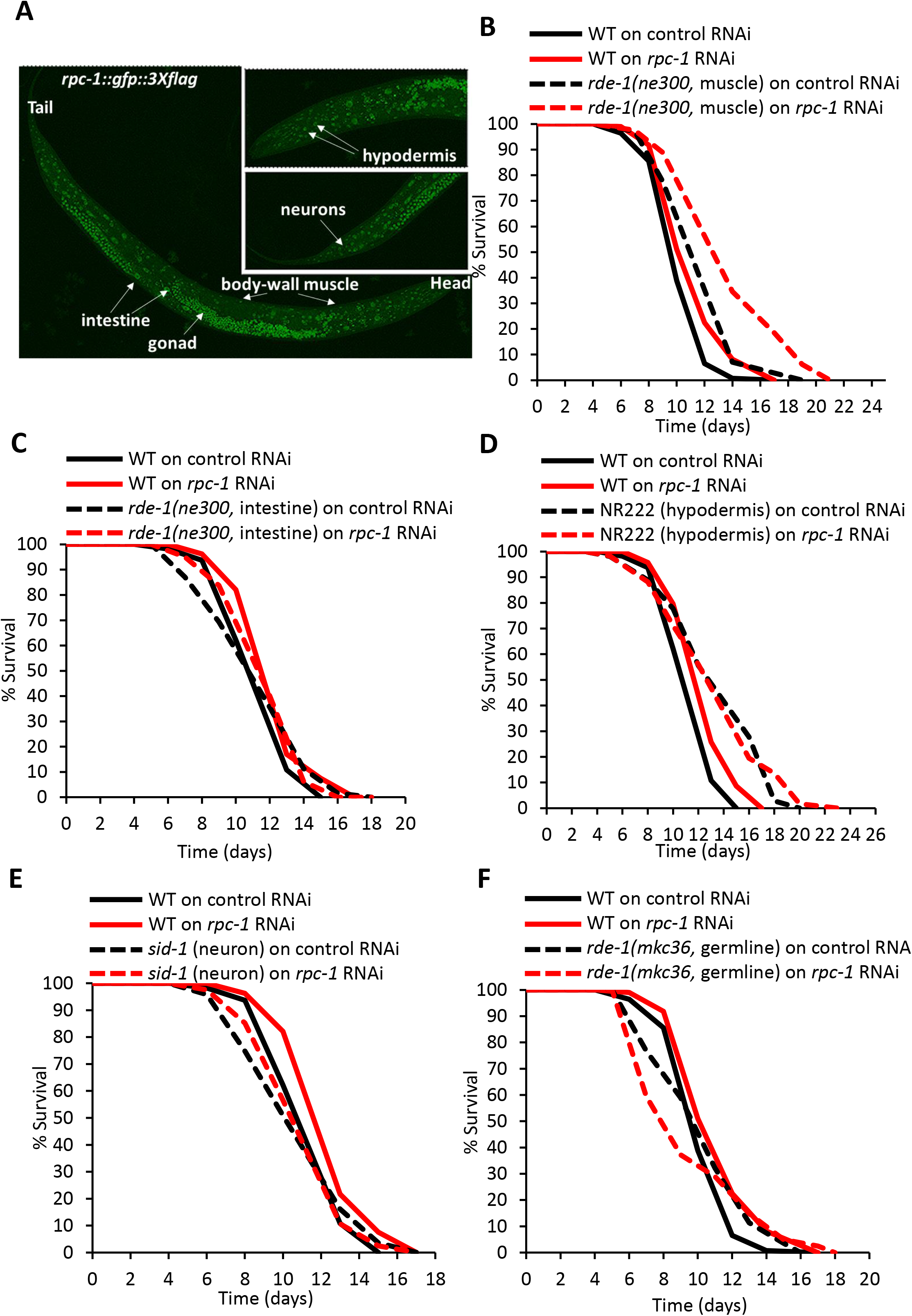
RPC-1 acts tissue specifically to control lifespan. **A)** RPC-1::GFP expression in L4 *C. elegans*. In each labelled tissue RPC-1::GFP is localized in the nuclei. Representative images shown. We also noted similar expression patterns in throughout adulthood (see also Fig. S3A). **B-F)** *rpc-1* RNAi specifically in body-wall muscle extends lifespan, but *rpc-1* knockdown in the intestine, hypodermis, neurons or germline does not extend lifespan. **B-F)** One representative experiment is shown, refer to Table 2 for data on all replicates and statistical analysis. Note: The result in **C)** contrasts with data using an alternative intestine-specific RNAi line in which *rpc-1* RNAi extends lifespan (see STAR methods and Fig. S3B).

To explore the requirement of these individual tissues for Pol III-knockdown mediated longevity, we utilized strains harboring a mutation in the dsRNA processing-pathway gene *rde-1*. The knockout mutant *rde-1(ne300)* is strongly resistant to dsRNA induced via the feeding RNAi method; with transgenic rescue of RDE-1 in different tissues allowing tissue-specific knockdown using RNAi^42^. We knocked-down *rpc-1* using RNAi from early adulthood in the muscle, intestine, hypodermis, neurons and germline, and measured their lifespan. Interestingly, *rpc-1* RNAi extended lifespan when knocked-down specifically in the body-wall muscles indicating a role for this tissue in mediating Pol III knockdown longevity (Fig. 3B, Table 2). However, we found that *rpc-1* RNAi in the hypodermis, germline, neurons or intestine could not increase lifespan compared to control (Fig. 3C-F, Table 2). Taken together, this suggests that Pol III acts in *C. elegans* muscle to inhibit lifespan under normal conditions.

**Table 2.**
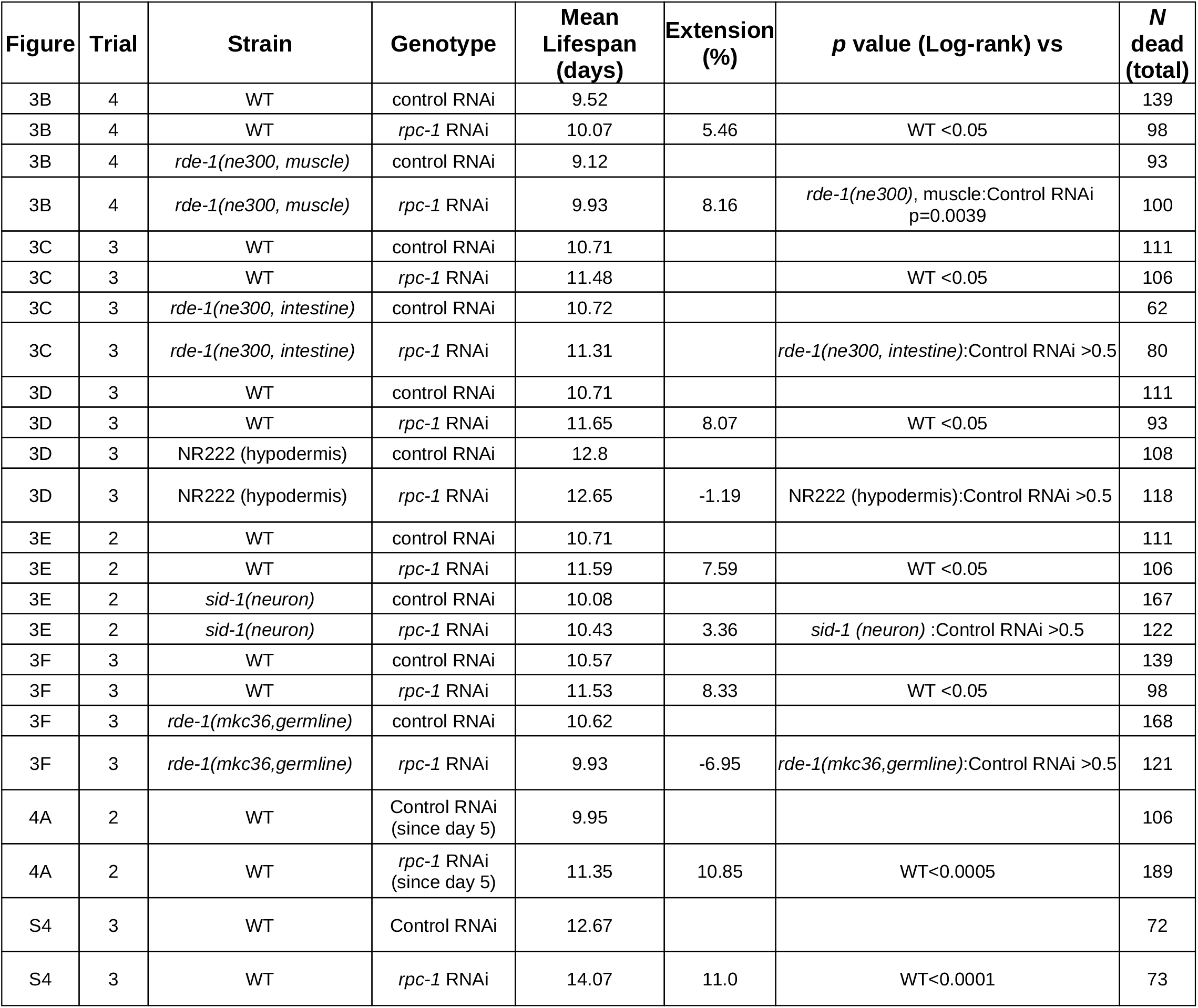

### Late-life knockdown of RNA Polymerase III is sufficient to extend lifespan and improve age-related health

In mice, the inhibition of TOR by rapamycin, has been shown to extend lifespan even when fed late in life^43^. Given that TOR - Pol III axis is important for adult lifespan (Fig. 2C, Table 1), we asked whether Pol III knockdown in late adulthood could also extend longevity in *C. elegans*. To test this, we raised worms on *E. coli* OP50-1 until day 5 of adulthood and then transferred them to either control or *rpc-1* RNAi plates. Excitingly, compared to control fed animals, worms on *rpc-1* RNAi lived longer, suggesting that Pol III knockdown in late adulthood effectively extends longevity (Fig. 4A, Table 2). Indeed, this late-life knockdown of Pol III resulted in a lifespan almost identical to that incurred by L4 RNAi treatment (Fig. S4, Table 2, Filer et al.^18^). Overall, these results show that Pol III knockdown in late adulthood is sufficient to extend *C. elegans* lifespan, strengthening our earlier finding that the beneficial longevity effects of Pol III are post reproduction.

**Figure. 4.**
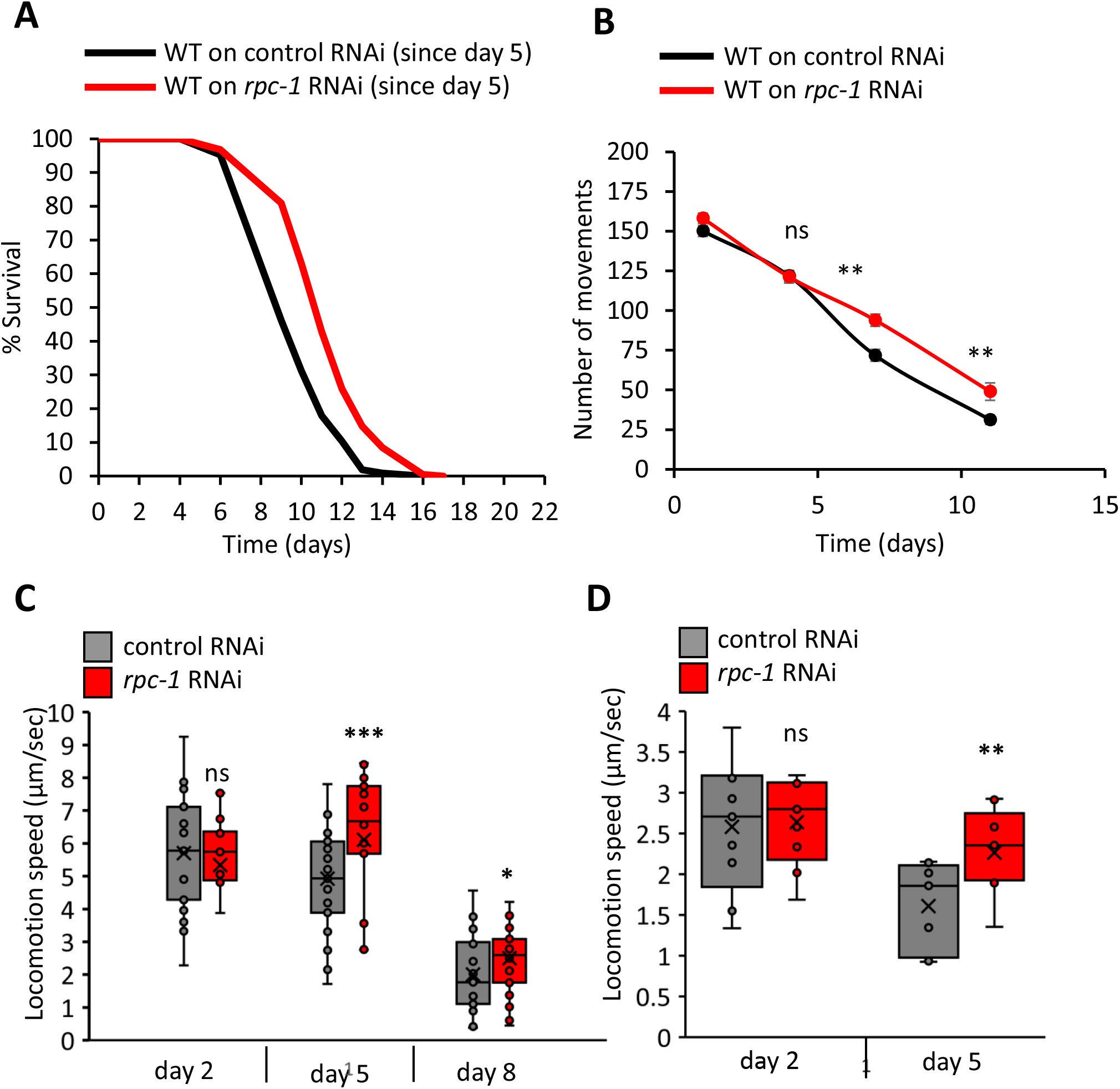
Reducing RNA polymerase III in late life extends lifespan and improves health. **A)** *rpc-1* RNAi in late-adulthood extends lifespan. One representative experiment is shown, refer to Table 2 for data on all replicates and statistical analysis. **B)** *rpc-1* RNAi improves *C. elegans* thrashing ability in liquid as they age. Measurements taken at days 1, 4, 7 and 11 of adulthood. Combined data from 3 trials shown, n=< 60 worms per group. *Student’s t-test* against age-matched control: **,*p*<0.005; ns, non-significant. **C)** *rpc-1* RNAi improves normal *C. elegans* movement in late life. **D)** *rpc-1* RNAi specifically in muscle improves normal *C. elegans* movement in late life. Note that the strain used for muscle specific RNAi also carries a *rol-6* marker, therefore the average movement is decreased compared to those exhibiting normal sinusoidal movement. In **C and D**) One representative experiment is shown. n=<20 worms per group. One-way ANNOVA: ***, *p*<0.0001; **,*p*<0.005; *,*p*<0.05; ns, non-significant.

Healthspan is an important indicator of youthfulness and virility and ideally, an increase in lifespan is complemented by improvement in organismal healthspan^44^. We investigated whether Pol III knockdown could also enhance the health and fitness of worms across their life by testing their ability to move in liquid media using a thrashing assay. As expected, thrashing ability decreased with age but compared to controls, worms on *rpc-1* RNAi moved better at later time points (day 7 and 11 of adulthood) than controls, suggesting improved fitness and better late-life health (Fig. 4B). We then compared the ability of *C. elegans* to move independently on food over their lifespan by videoing their natural movement on *E. coli* seeded plates. Similarly to the trashing assay *C. elegans* moved around on plates less as they aged, but older worms treated with *rpc-1* RNAi moved better compared to controls (Fig. 4C, day 5 adults *p*= 0.0003, day 8 adults *p*=0.002 compared to age-matched controls). Together with the thrashing data this supports a role for Pol III in mediating life-long health as well as overall lifespan.

Given that it is Pol III knockdown in muscle is sufficient to promote longevity, we also tested whether muscle-specific *rpc-1* RNAi improved age-related fitness. We found that indeed *rpc-1* knockdown in the muscle was sufficient to improve overall body movement on day 5 of adulthood (Fig. 4D, *p*= 0.001 compared to control). This supports our finding that *rpc-1* is acting in muscle to control age-related health. Taken together, these results suggest that Pol III activity is required for larval growth and development in *C. elegans*, and knockdown of Pol III in adulthood lead to significant increase in lifespan and healthspan.

## Discussion

### RNA Polymerase III and TOR signalling interact temporally to control development and lifespan

Both TOR signalling and RNA Polymerase III are important regulators of anabolic processes, and control an wide array of cellular functions including reproduction and lifespan^16^. Here we show that TOR and Pol III interact in a time dependent manner, and that knockdown of each component individually, sequentially or in parallel influences the outcome of both reproductive and longevity processes (Fig. 5). In *C. elegans*, we manipulated TOR signalling using both genetic mutants (that constitutively reduce TOR signalling throughout life) and TOR RNAi (which reduces TOR signalling the late L4 stage). We show that the genetic relationship between TOR signaling and Pol III is timing dependent. The requirement and collaboration of TOR and Pol III to support developmental processes, and the fact that their reduction in adulthood extends lifespan supports the antagonistic pleiotropy theory of ageing; whereby certain genes are switched ‘on’ to promote the anabolic programs leading to growth, sexual maturity and reproductive fitness, and ‘hyperfunction’ of these genes in adulthood leads to senescence and ageing^45^.

**Figure. 5.**
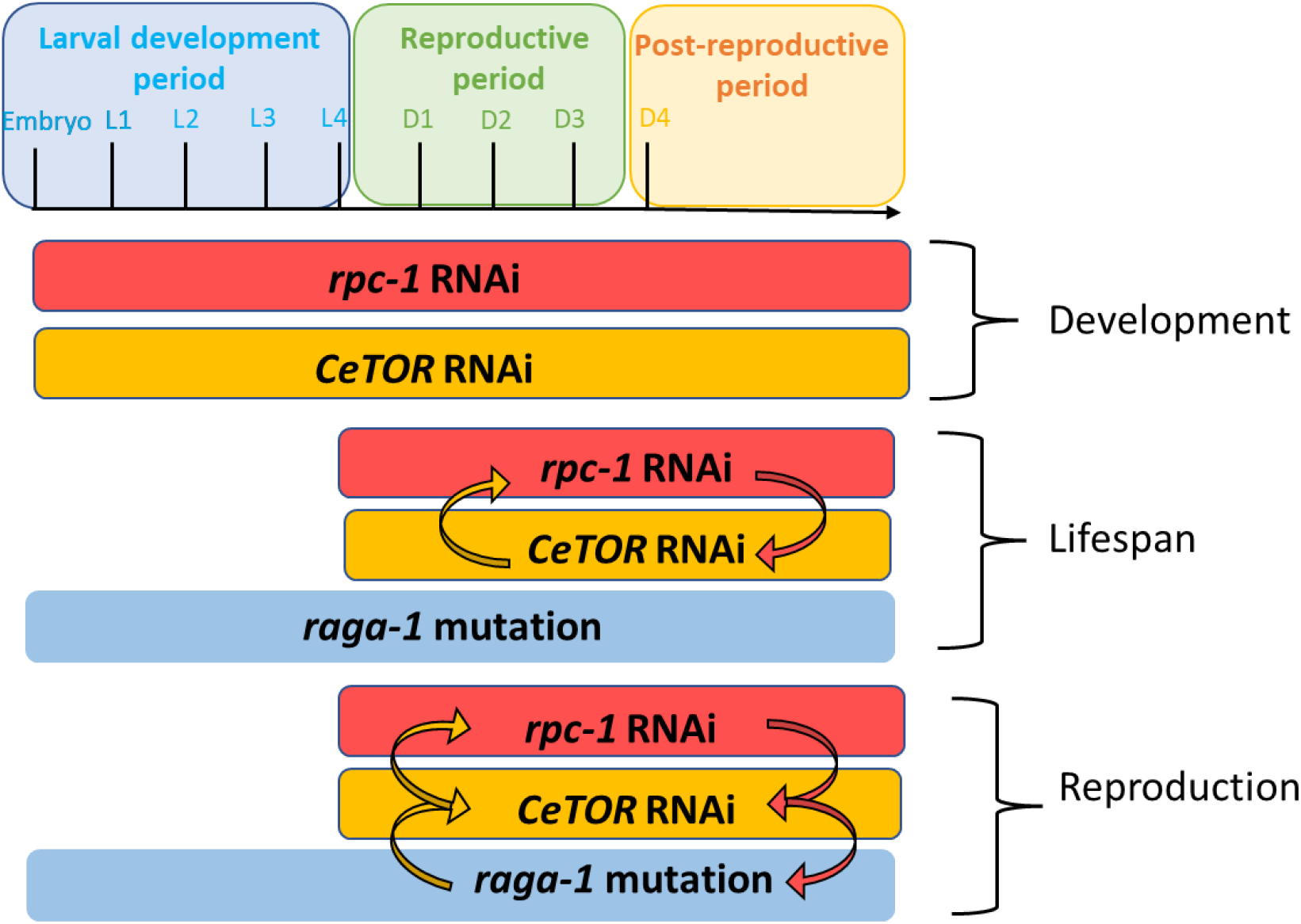
Temporal genetic interactions between TORC1 and RNA Polymerase III. Model summarizing the interactions between TORC1 and RNA Polymerase III required for development, reproduction and lifespan. This relationship is highly temporal: Reducing TOR signalling specifically during adulthood requires RNA Pol III for reproduction and lifespan, whereas TORC1 reduction during the embryonic and larval period requires other mediators to promote reproduction and longevity.

The contrasting effects on fitness and longevity as observed by knockdown of Pol III in early development as opposed to late adulthood are similar to those observed for other signaling pathways including mTOR and insulin. Knockdown of *daf-2* from early to mid-adulthood also promotes longevity while its knockdown from early larval stages reduces fecundity and reproductive fitness^46^. Late-life interventions targeting IGF-1 receptor have also been shown to improve lifespan and healthspan in mice^47^. However, whilst the lifespan of *daf-2* worms cannot be further extended by *CeTOR* inhibition, suggesting a common mechanistic pathway^28^, we did not find evidence for a genetic interaction with the *C. elegans* insulin receptor *daf-2* (Fig. S2A). In fact, *daf-2* mutation and *rpc-1* RNAi were additive in our hands implying independent downstream effectors of Pol III and DAF-2. Our similar results with germline-signaling mutants (Fig. S2B and C) which increase longevity by reducing germline activity lead us to speculate that this could be in part due to the common transcriptional denominator DAF-16, which integrates signals both from the IGF-1/ DAF-2 axis and germline-loss driven longevity^48–50^, while Pol III knockdown bypasses DAF-16 requirements entirely.

### RNA Polymerase III inhibition as a modulator of late-life health and lifespan

Studies in a wide variety of model organisms show that mTOR inhibition increases longevity and improves organismal health, spurring interest in mTOR inhibitors for slowing down human ageing^51^. However, this can occur at the cost of development e.g. treating female flies with rapamycin increased lifespan, but significantly reduced brood size^52,54^. In addition, clinical studies have shown that chronic exposure to mTOR inhibitors can result in serious side effects including immunosuppression and metabolic dysfunction, making them unattractive anti-ageing therapies^27^. Consequently transient knockdown of mTOR genetically, or by inhibitors, is a fast emerging paradigm for targeting mTOR complexes. Studies targeting mTORC1 by rapamycin even late in life, have shown to extend lifespan in mice models, without major side-effects^43,53^. Here we show that while Pol III knockdown during early larval stages leads to developmental abnormalities and larval arrest (Fig. 1B), Pol III knockdown post-reproduction on the fifth day of adulthood, extends lifespan suggesting that similarly to TOR, even late perturbations in Pol III activity are sufficient to promote longevity (Fig. 4A). Importantly, we also show that Pol III inhibition is sufficient to improve age-related decline in movement (Fig. 4D). Overall, our results make RNA Pol III a promising target molecule for ameliorating post-reproductive age-related decline.

### RNA Polymerase III acts in muscle to drive ageing

It is known that aging progresses heterogeneously and various tissue-specific responses have been identified to perturb organismal aging. Earlier studies have shown that knockdown of Pol III in fly gut improves gut barrier function and is sufficient to ameliorate age-related impairments in gut health which may ultimately promote longevity ^18^. In *C. elegans*, multiple studies have shown tissue-specific requirements of longevity genes^50,55^ and Pol III RNAi mediated longevity can also be mediated via the intestine^18^. The broad expression of *rpc-1::GFP* (Fig. 3A and Fig. S3A), led us to explore its longevity and health functions in other tissues and found that muscle-specific knockdown of *rpc-1* was sufficient to extend lifespan (Fig. 3B). Multiple studies, in both worms and mammals, have shown that improving skeletal muscle function can improve overall health and longevity^56–58^. In *C. elegans* even reducing translation in body-wall muscle during development shortens lifespan and increases reproduction^59^. Whilst loss of RAGA-1 in the nervous system increases lifespan via maintaining mitochondrial fusion in the muscles^60^. One possible mechanism for this, is the improvement in skeletal musculature of *C. elegans*. Indeed, we also saw an improvement in age-related movement/fitness in this context (Fig. 4D).

Lowering translation or mTORC1 activity in adulthood consistently increases lifespan^20^, reduces brood size and improves motility^31,61^. Surprisingly though, suppressing TORC1 specifically in body-wall muscle increased reproduction and slowed motility, indicating a role for the muscle in TOR-mediated activity^62^. We could not test if muscle-specific reduction of Pol III during development altered lifespan owing to the strong larval arrest phenotype of *rpc-1* RNAi but suspect that such an intervention may perturb global protein translation and remodel inter-tissue communication via one or many of the small-RNAs transcribed by Pol III.

### Exploring additional mediators of the Pol III response

It has been well documented that inhibition of translation delays ageing and evidence suggests that reducing global translation can maintain protein homeostasis, dysregulation of which leads to aging and senescence^61,63,64^. It is likely that Pol III reduction leads to a decrease of two major RNA types, tRNAs and 5s rRNA, subsequently downregulating global protein translation. Hence we were surprised to find that *rpc-1* RNAi induced longevity was additive with *C. elegans* mutants of other key translation mediators (Fig. S1). This implies a more nuanced role of Pol III in maintaining a healthy proteome, possibly via improving protein folding or reducing insoluble protein loads, which would be important for lifespan and age-related disease. In the future, addressing the impact of Pol III inhibition on protein levels and quality should prove interesting.

## Acknowledgements

This work was funded by a BBSRC NI award BB/R003629/1 to JMAT and BBSRC award BB/S014365/1 to JMAT, NA and CS. Some strains were obtained from *Caenorhabditis* Genetics Center, Minnesota, U.S.A, which is funded by the NIH Office of Research Infrastructure Programs (P40 OD01044). *ppp-1(syb7781)* was a kind gift from the Denzel lab (Cologne, Germany).

## Author contributions

Conceptualization, JMAT; Methodology, JMAT, YM, IGS, RRD; Investigation, JMAT, YM, IGS, RRD; Writing Original Draft, JMAT, YM; Writing Review & Editing, JMAT, NA, CS; Funding Acquisition, JMAT, NA, CS; Supervision, JMAT.

## Data Availability Statement

All relevant data are within the manuscript and its Supporting Information files.

## Materials and Methods

### *C. elegans* maintenance and strains

*C. elegans* were routinely grown and maintained on Nematode Growth Media (NGM) seeded with *Escherichia coli* OP50-1. The wild-type strain was Bristol N2. Some strains were provided by the CGC, which is funded by NIH Office of Research Infrastructure Programs (P40 OD010440). The following strains were used:VC222 *raga-1(ok386)*, RB1206 *rsks-1(ok1255)*, RB579 *ife-2 (ok306), ppp-1(syb7781)*, DR1567 *daf-2*, CB4037 *glp-1(e2141)*, SS104 *glp-4(bn2)*, IG1839 *frSi17 II; frIs7 IV; rde-1(ne300) V* (intestine-specific RNAi), DCL569 *mkcSi13 II; rde-1(mkc36) V* (germline-specific RNAi). WM118 *rde-1(ne300) V; neIs9 X* (muscle-specific RNAi). TU3401 *sid-1(pk3321) V; uIs69 V* (neuronal-specific RNAi), NR222 *rde-1(ne219) V; kzIs9* (hypodermis-specific RNAi), VP303 *rde-1(ne219) V; kbIs7* (intestine-specific RNAi).

### Lifespan assay

Longevity assays were performed as previously described (Filer et al, 2017). Briefly, synchronous L1 animals were placed on NGM plates seeded with *E. coli* OP50-1 until they reached L4. From L1 to L4 larval stage animals were maintained at 20°C. At L4 stage, worms were shifted to plates seeded with *E. coli* HT1115 carrying appropriate RNAi clones. These RNAi clones were obtained from the Ahringer RNAi library. The plates were supplemented with 50 μM FuDR solution and lifespans were carried out at 25°C. For combinatorial RNAi knockdown, the two respective RNAi cultures were mixed 1:1 before seeding the plates. In these experiments, controls were also diluted 1:1 with HT115 expressing empty pL4440. Live, dead and censored worms were calculated every 2–3 days in the worm populations by scoring movement with gentle prodding when necessary. Data were analyzed and statistics performed using the online freeware OASIS (http://sbi.postech.ac.kr/oasis/surv/).

### Brood size measurements

Animals were grown on *E. coli* OP50-1 plates at 20°C till L4 larval stage. At L4, approximately 15 animals were placed on individual NGM plates seeded with the appropriate RNAi. For combinatorial RNAi, see protocol in lifespan section. Progeny production was assessed at 25°C and each parent animal moved to a fresh plate daily until egg-laying ceased. The total number of eggs and larvae were counted after 24 hrs. Plates in which the animals went missing or showed intestinal bursting were not included in the final analysis.

### Thrashing Assay

Animals were raised to L4 as for lifespan assays. At L4, the animals were moved to control RNAi or *rpc-1* RNAi seeded plates. On days 4, 7 and 11 of adulthood worms were put in the isotonic M9 buffer and allowed to acclimatise for one minute before measuring the number of side-to-side mid-body movements per minute. Fifteen animals were analysed per condition.

### Microscopy

*C. elegans* carrying the *rpc-1::gfp::3Xflag* reporter were grown in fed conditions to the L4 stage. For imaging, animals were immobilized on 2% agarose pads using 0.06% Levamisole and imaged immediately using a Zeiss Confocal Ultra at 40X zoom.

### Speed measurement

*C. elegans* were raised and maintained as for the lifespan assays. At each timepoints 20-30 worms were moved to a thinly seeded 60 mm plate and were allowed to acclimatise for 10 minutes. Their individual movements were then videoed and recorded using Wormtracker (MBF Bioscience) for 2 minutes with 60 fpm settings. Average speeds for individual animals were calculated using MS Excel.

## Figure legends

**Figure S1.**
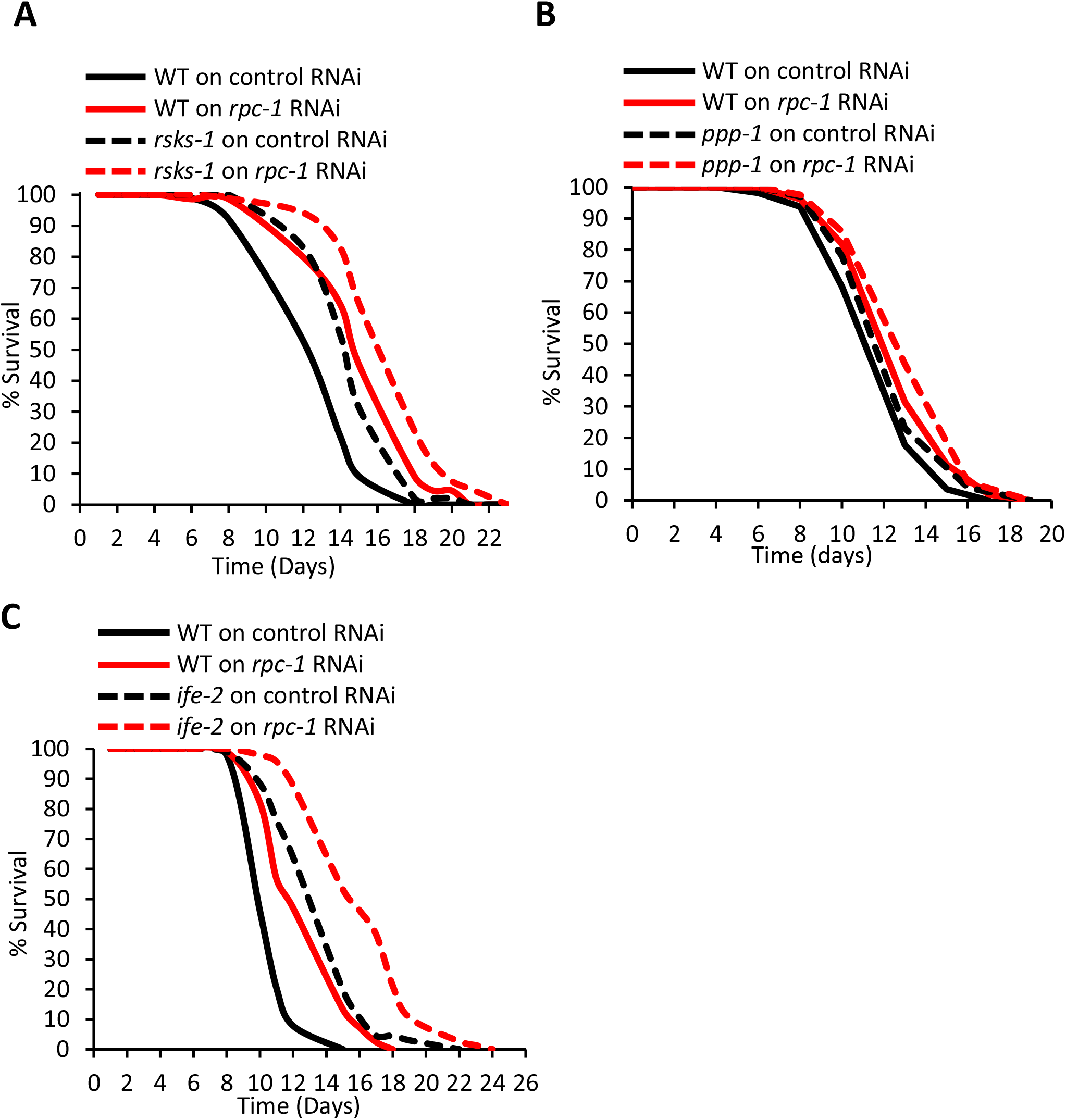
RNA polymerase III does not genetically interact with global translation modulators. **A-C)** *rpc-1* RNAi additively increases lifespan with *rsks-1(ok1255), ppp-1 (syb7781) and ife-2(ok306)* mutants. One representative experiment is shown, refer to Table S1 for data on all replicates and statistical analysis.

**Figure S2.**
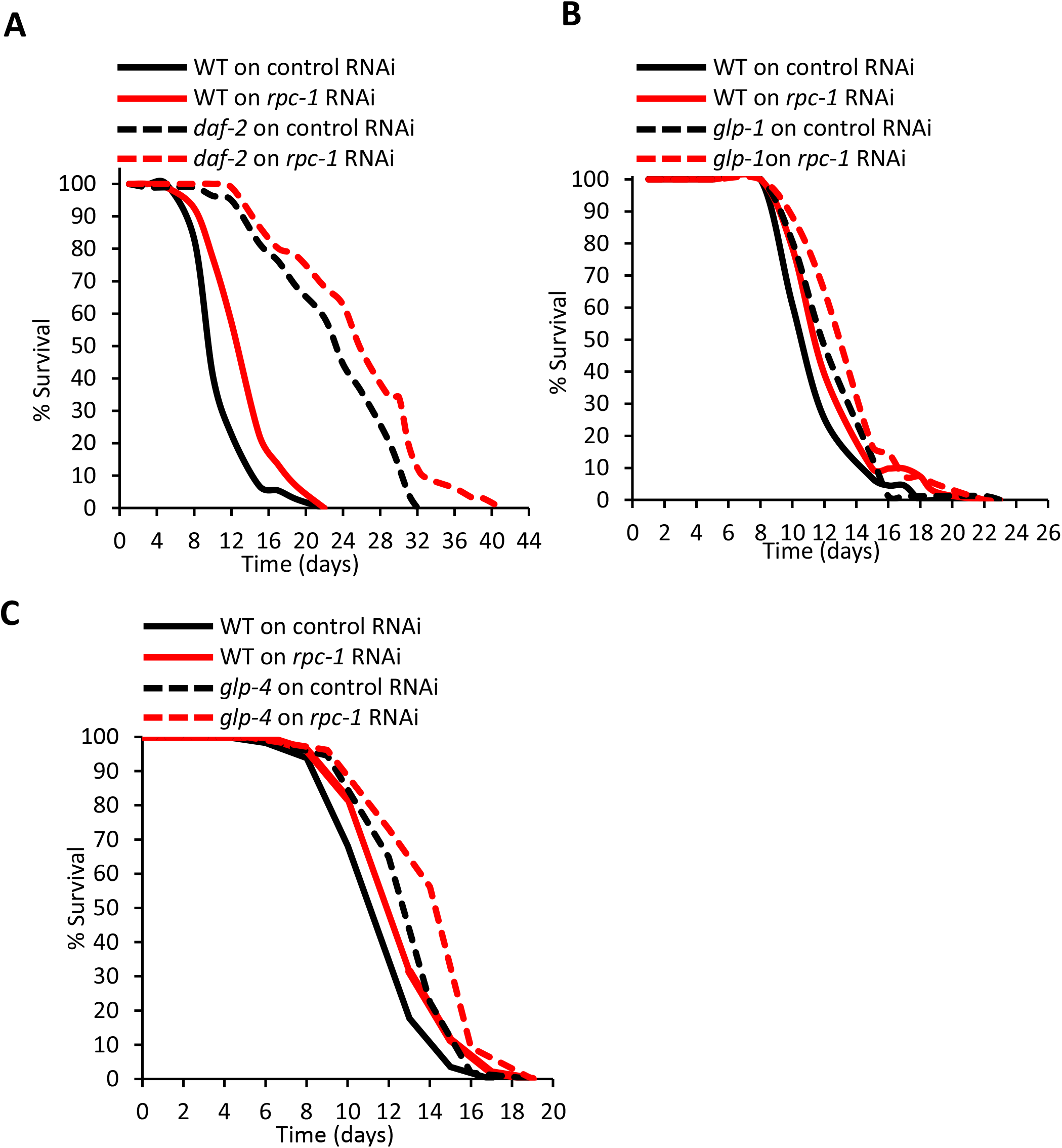
RNA polymerase III does not genetically interact with insulin or germline signalling. **A-C)** *rpc-1* RNAi additively increases lifespan with *daf-2(m577), glp-1(e2141)* and *glp-4(bn2)* mutants. One representative experiment is shown, refer to Table S2 for data on all replicates and statistical analysis.

**Figure S3.**
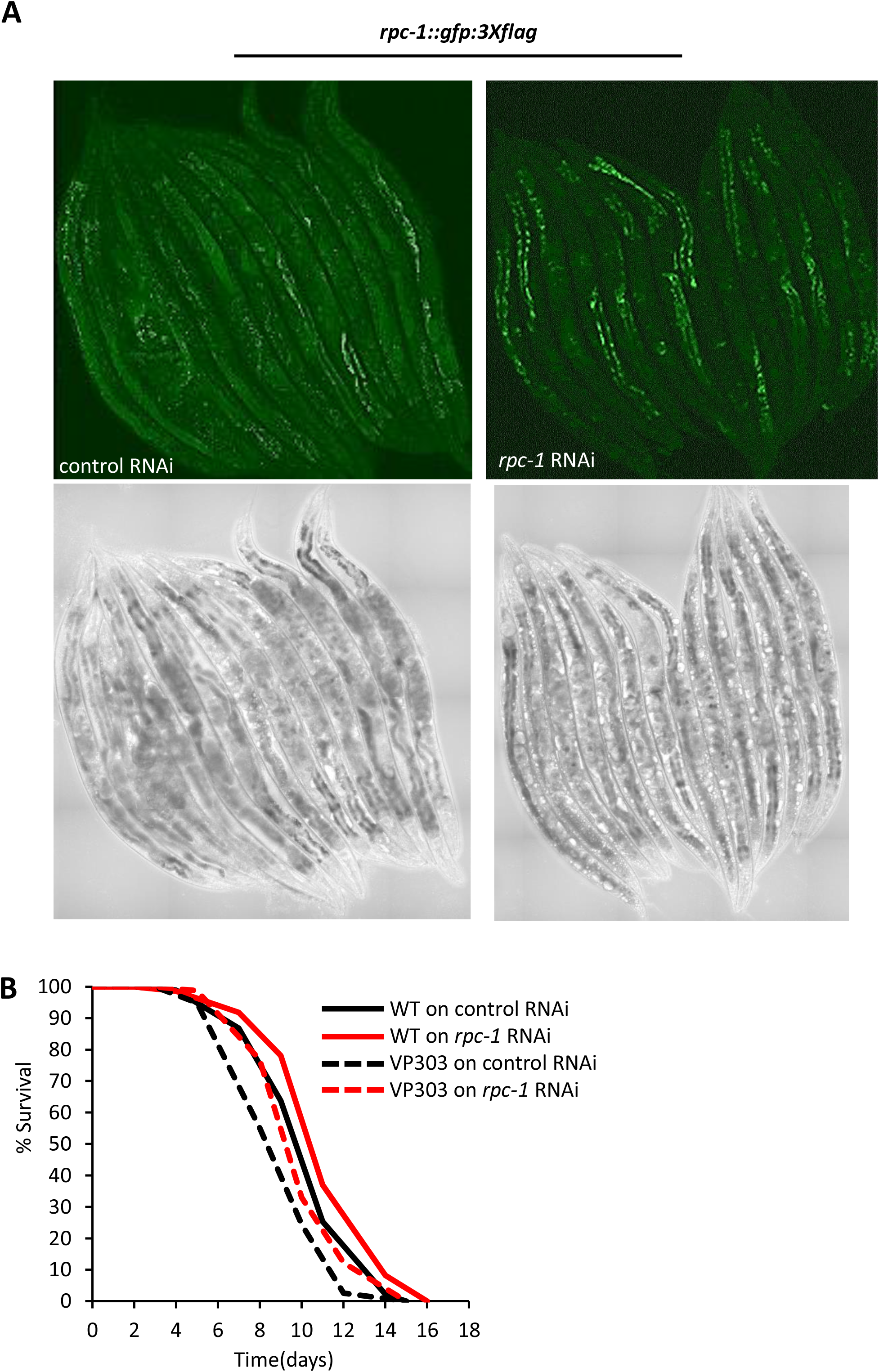
Tissue specific expression and function of RNA Polymerase III. **A)** RPC-1::GFP expression in *C. elegans* fed either control and *rpc-1* RNAi in day 5 adults. RPC-1::GFP expression is reduced on worms fed with *rpc-1* RNAi. Bacteria expressing RNAi was fed for 5 days prior to imaging. **B)** *rpc-1* RNAi in the intestine-specific RNAi strain VP303 significantly extends lifespan as reported in Filer et al., 2017. One representative experiment is shown, refer to Table S1 for data on all replicates and statistical analysis. This observation is in contrast to our published data wherein *rpc-1* RNAi extends the lifespan of a different intestine-specific RNAi strain *rde-1(ne219)*. However, this difference is likely attributed to the allele-specific differences between *rde-1(ne219)* and *rde-1(ne300)*; *ne219* retains significant RNAi processing capacity and is not completely null whereas *ne300* deletion allele is completely resistant to RNAi (Watts et al.^42^). Thus, the *rde-1(ne300)* strain we initially tested is likely to be ‘leaky’ and RNAi knockdown might not be limited to the gut of the worms.

**Figure S4.**
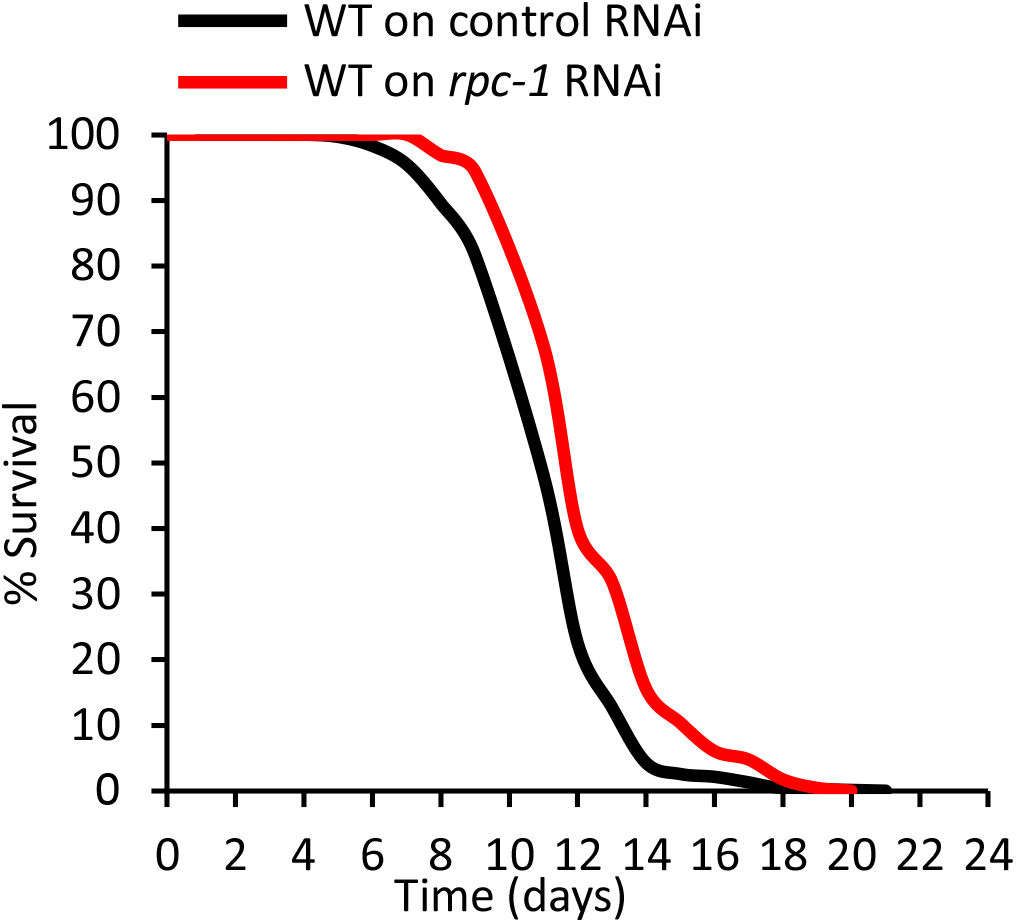
*rpc-1* RNAi from L4 larval stage extends lifespan to a similar extent as late-life *rpc-1* RNAi. *rpc-1* RNAi from L4 larval stage extends lifespan similarly to RNAi induced on day 5 adulthood (Fig 4A). One representative experiment is shown, refer to Table 2 for data on all replicates and statistical analysis.

**Table S1.**
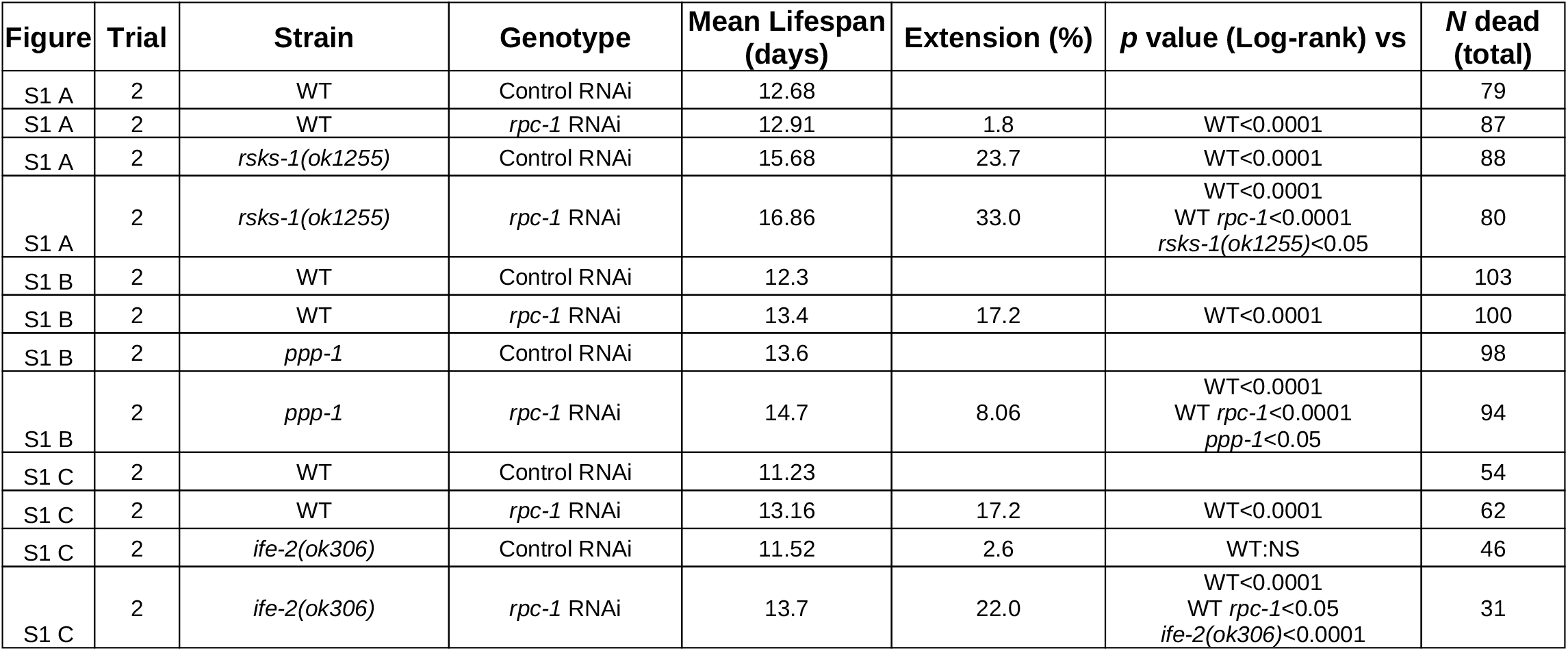

**Table S2.**
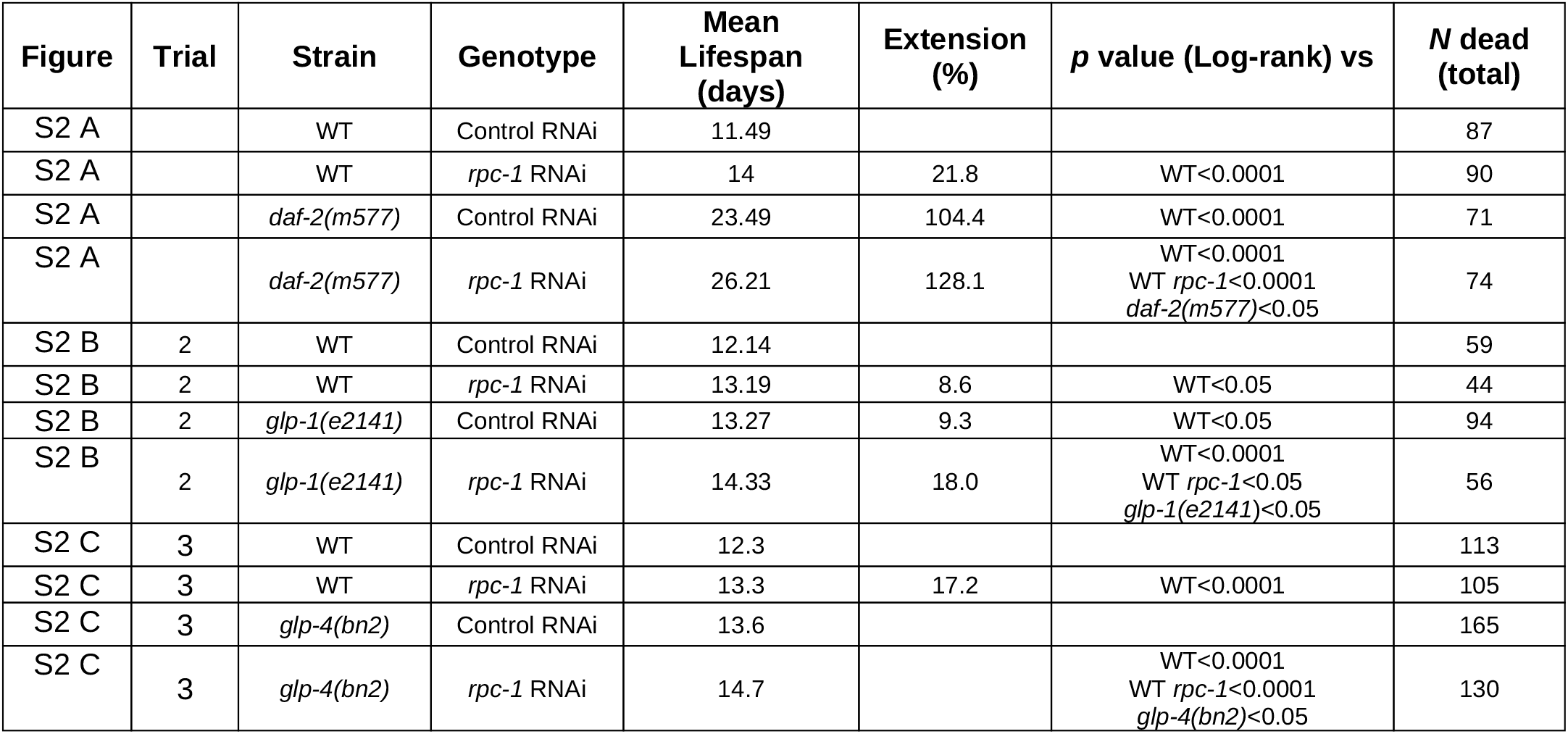

| Figure | Trial | Strain | Genotype | Mean Lifespan (days) | Extension (%) | p value (Log-rank) vs | N dead (total) |
| --- | --- | --- | --- | --- | --- | --- | --- |
| S2 A |  | WT | Control RNAi | 11.49 |  |  | 87 |
| S2 A |  | WT | *rpc-1* RNAi | 14 | 21.8 | WT<0.0001 | 90 |
| S2 A |  | *daf-2(m577)* | Control RNAi | 23.49 | 104.4 | WT<0.0001 | 71 |
| S2 A |  | *daf-2(m577)* | *rpc-1* RNAi | 26.21 | 128.1 | WT<0.0001<br>WT *rpc-1*<0.0001<br>*daf-2(m577)*<0.05 | 74 |
| S2 B | 2 | WT | Control RNAi | 12.14 |  |  | 59 |
| S2 B | 2 | WT | *rpc-1* RNAi | 13.19 | 8.6 | WT<0.05 | 44 |
| S2 B | 2 | *glp-1(e2141)* | Control RNAi | 13.27 | 9.3 | WT<0.05 | 94 |
| S2 B | 2 | *glp-1(e2141)* | *rpc-1* RNAi | 14.33 | 18.0 | WT<0.0001<br>WT *rpc-1*<0.05<br>*glp-1(e2141)*<0.05 | 56 |
| S2 C | 3 | WT | Control RNAi | 12.3 |  |  | 113 |
| S2 C | 3 | WT | *rpc-1* RNAi | 13.3 | 17.2 | WT<0.0001 | 105 |
| S2 C | 3 | *glp-4(bn2)* | Control RNAi | 13.6 |  |  | 165 |
| S2 C | 3 | *glp-4(bn2)* | *rpc-1* RNAi | 14.7 |  | WT<0.0001<br>WT *rpc-1*<0.0001<br>*glp-4(bn2)*<0.05 | 130 |

